# Altered Functional Network Energy Across Multiscale Brain Networks in Preterm vs. Full-Term Subjects: Insights from the Adolescent Brain Cognitive Development (ABCD) Study

**DOI:** 10.1101/2025.02.09.637316

**Authors:** Qiang Li, Dawn Jensen, Zening Fu, Teddy Jakim, Masoud Seraji, Selim Süleymanoğlu, G. Hari Surya Bharadwaj, Jiayu Chen, Vince D. Calhoun, Jingyu Liu

**Author notes:** 47th IEEE Engineering in Medicine and Biology Society (EMBC 2025).

## Abstract

Infants born prematurely, or preterm, can experience altered brain connectivity, due in part to incomplete brain development at the time of parturition. Research has also shown structural and functional differences in the brain that persist in these individuals as they enter adolescence when compared to peers who were fully mature at birth. In this study, we examined functional network energy across multiscale functional connectivity in approximately 4600 adolescents from the Adolescent Brain Cognitive Development (ABCD) study who were either preterm or full term at birth. We identified three key brain networks that show significant differences in network energy between preterm and full-term subjects. These networks include the visual network (comprising the occipitotemporal and occipital subnetworks), the sensorimotor network, and the high cognitive network (including the temporoparietal and frontal subnetworks). Additionally, it was demonstrated that full-term subjects exhibit greater instability, leading to more dynamic reconfiguration of functional brain information and increased flexibility across the three identified canonical brain networks compared to preterm subjects. In contrast, those born prematurely show more stable networks but less dynamic and flexible organization of functional brain information within these key canonical networks. In summary, measuring multiscale functional network energy offered insights into the stability of canonical brain networks associated with subjects born prematurely. These findings enhance our understanding of how early birth impacts brain development.

## I. Introduction

Babies born before full-term (usually before 40 weeks of gestational age) are classified as preterm [1]. Both internal factors, such as biological attributes, and external factors, like environmental conditions, can influence the likelihood of preterm birth [2]. The impact of preterm birth could be profound, including, but not limited to, disruptions in the structural and functional development of the brain, increased risk to neurodevelopmental disorders [3], [4], although the exact mechanisms remain unclear.

Numerous investigations have focused on examining abnormalities in brain structures and functions of preterm brain, specifically in white matter and gray matter [5], [6], [7]. Additionally, studies have aimed to characterize these alterations at the brain connectome level from both functional and structural perspectives [7], [8]. fMRI studies, in particular, have assessed the functional properties of the preterm brain, revealing reduced functional connectivity and more localized neural networks [6]. The preterm brain is more susceptible to developmental challenges, with higher rates of impairments in sensory, motor, and cognitive functions [6]. Furthermore, when analyzing resting-state network complexity, studies have shown that brain organization tends to become more ordered with age, though earlier birth is associated with greater chaos in brain activity [9]. Notably, the motor and sensory networks show the most significant increase in complexity and order as age increases [9], [10].

Although many studies have investigated differences between preterm and full-term brain, less is known about the potential long-term effect on the brain when children enter adolescence. Most studies have relied on small sample sizes and have focused mainly on structural brain development. The limited functional brain studies have focused on functional connectivity differences based on anatomical atlases and overlooked multiscale functional brain networks and network properties beyond pairwise correlation [11], [12].

In this study, we aim to address these limitations by analyzing brain functional networks in a large sample of children at age of 9-10 year old, by using a comprehensive data-driven independent component analysis (ICA) approach to identify multiscale canonical functional networks, and by assessing the energy of network, a network stability measure. Our focus is on examining long term effects on functional brain networks induced by preterm birth. This approach offers a novel way to explore brain functional organization both at the global whole brain level and at subnetworks within specific canonical brain networks.

## II. Methodology

### A. Participants with Preterm and Full-Term Birth

The data used in this study were obtained from the Adolescent Brain Cognitive Development (ABCD) Study (https://abcdstudy.org), hosted in the NIMH Data Archive (https://nda.nih.gov/). This multisite, longitudinal study aims to recruit over 10,000 children aged 9-10 and follow them for 10 years into early adulthood at 21 sites across the United States [13]. For the current analyses, we further selected participants based on parent report of length of pregnancy, with parturition at less than 40 weeks considered premature. Preterm subjects range in prematureness from one to ten weeks. Preterm and full-term subject groups were matched by sex as well as social economic status and were all nine years old at the time of their baseline scans. After preprocessing and performing quality control (QC) to select scans for further analysis, we included a total of 1,849 preterm subjects and 2,765 full-term subjects.

### B. rsfMRI Dataset Acquisition and Processing

The resting-state data was collected using three 3T scanner platforms, including Siemens Prisma, Philips, and GE 750 3T scanners. The imaging protocols are similar across scanners [13], which are listed below: repetition time (TR)/echo time (TE)=800/30ms, voxel spacing size = 2.4 × 2.4 × 2.4mm, number of slices = 60, flip angle (FA)=52°, field of view (FOV)=216 × 216mm, %FOV phase = 100%, and multiband acceleration=6. We preprocessed the raw data using a standard preprocessing pipeline that combines the FMRIB Software Library (FSL) v6.0 and Statistical Parametric Mapping (SPM) 12.

### C. Pipeline for Building Multiscale Functional Network Connectivity

In this study, we utilized the NeuroMark_fMRI_2.2 template (https://trendscenter.org/data/), a multiscale brain network template derived from over 100K subjects [14], [15]. These templates were derived from over 20 different datasets and processed using a group multi-scale ICA approach with 8 distinct model orders to achieve varying spatial resolutions [14]. By incorporating components across multiple scales, we can model a broader range of intrinsic connectivity networks (ICNs).

Finally, a total of 105 ICNs were identified and organized into seven major functional networks: visual network (VI, 12 subnetworks; occipitotemporal subnetwork (OT) and occipital subnetwork (OC)), cerebellar network (CB, 13 sub-networks), sub-cortical network (SC, 18 sub-networks; extended hippocampal subnetwork (EH), extended thalamic subnetwork (ET), and basal ganglia subnetwork (BG)), sensorimotor network (SM, 14 sub-networks), high cognition network (HC, 22 sub-networks; insular-temporal subnetwork (IT), temporoparietal subnetwork (TP), and frontal subnetwork (FR)), triple network network (15 sub-networks; central executive subnetwork (CE), default mode subnetwork (DM), and salience subnetwork (SA)), and paralimbic network (PL, 11 sub-networks), as illustrated in ***Fig.1A***.

We then applied the Multivariate Objective Optimization ICA with Reference (MOO-ICAR) framework [16] for spatially constrained independent component analysis (scICA) on the selected preprocessed preterm and full-term subjects scans from the ABCD study. This allowed us to extract subject-specific ICNs and their corresponding time courses. Multiscale functional connectivity was then estimated using Pearson correlation for each subject in both groups.

### D. Estimating Functional Network Energy

Assessing brain network energy through structural balance theory offers insights into the organization of functional connections [17], [18]. Low energy indicates stability, with well-coordinated connections that reduce internal conflicts and promote efficient information processing, but it also reduces flexibility and the capacity for more dynamic reorganization of information. High energy, indicative of instability, reflects tension and conflicting connections, enabling dynamic reconfiguration of information organization. These energy dynamics emphasize the brain’s capacity to balance stability and flexibility. The functional network energy [18] can be formulated as,

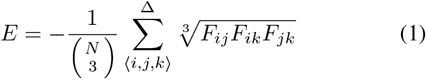

where *F*_*ij*_, *F*_*ik*_, and *F*_*jk*_ represent the functional connection weights between ICNs *i* and *j, i* and *k*, and *j* and *k*, respectively. *N* = 105 is the number of ICNs, and 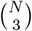 represents the number of triangles in a fully connected network. The combination term normalizes the energy between -1 and 1 by dividing by the number of triangles.

The summation is performed across all possible triangles formed by regions *i, j*, and *k*, as indicated by the triangle symbol above the summation. *F*_*ij*_, *F*_*ik*_, and *F*_*jk*_ represent the weighted connections of the triangle. Triadic interactions offer a new approach to exploring beyond pairwise functional connections, providing us with additional valuable information [18], [19], [20], [21]. If the product is positive, the triangle is balanced. If the product is negative, the triangle is imbalanced [22]. Summing across all triangles helps evaluate network stability. Balanced triangles promote stability, while imbalanced triangles lead to instability. Given the negative sign in the equation, a connectome with mostly balanced (stable) triangles and well-coordinated connections has negative energy. Conversely, a connectome dominated by imbalanced (unstable) triangles and conflicting connections has positive energy [17], [18], [22].

### E. Permutation Statistical Test on Network Energy

To identify the significant differences of network energy between the two groups, we first computed the mean difference of network energy between the two groups (preterm vs. full-term) as the observed value. Next, we performed 10,000 permutations and tests on network energy values. For each permutation, we combined the functional network energy from both groups into a single dataset, then randomly shuffled and split back into two groups, preserving the original group sizes. The test statistic, which is the mean difference in network energy between the two groups, was calculated for each permuted dataset. Finally, we determined the p-value as the proportion of permuted test statistics that were greater than or equal to the absolute value of the observed test statistic, thereby evaluating the significance of the observed difference between the two groups. We performed a permutation test on the whole-brain network and fourteen networks.

## III. Results

### A. Multiscale Functional Connectivity in Preterm versus Full-Term Subjects

A total of 105 ICNs and their corresponding time series were identified across the entire brain for each subject in both the preterm and full-term groups. The average preterm functional connectivity matrix from 1,849 subjects and the average full-term functional connectivity matrix from 2,765 subjects were computed, along with their difference functional connectivity matrix, as presented in ***Fig. 1B***.

**Fig. 1:**
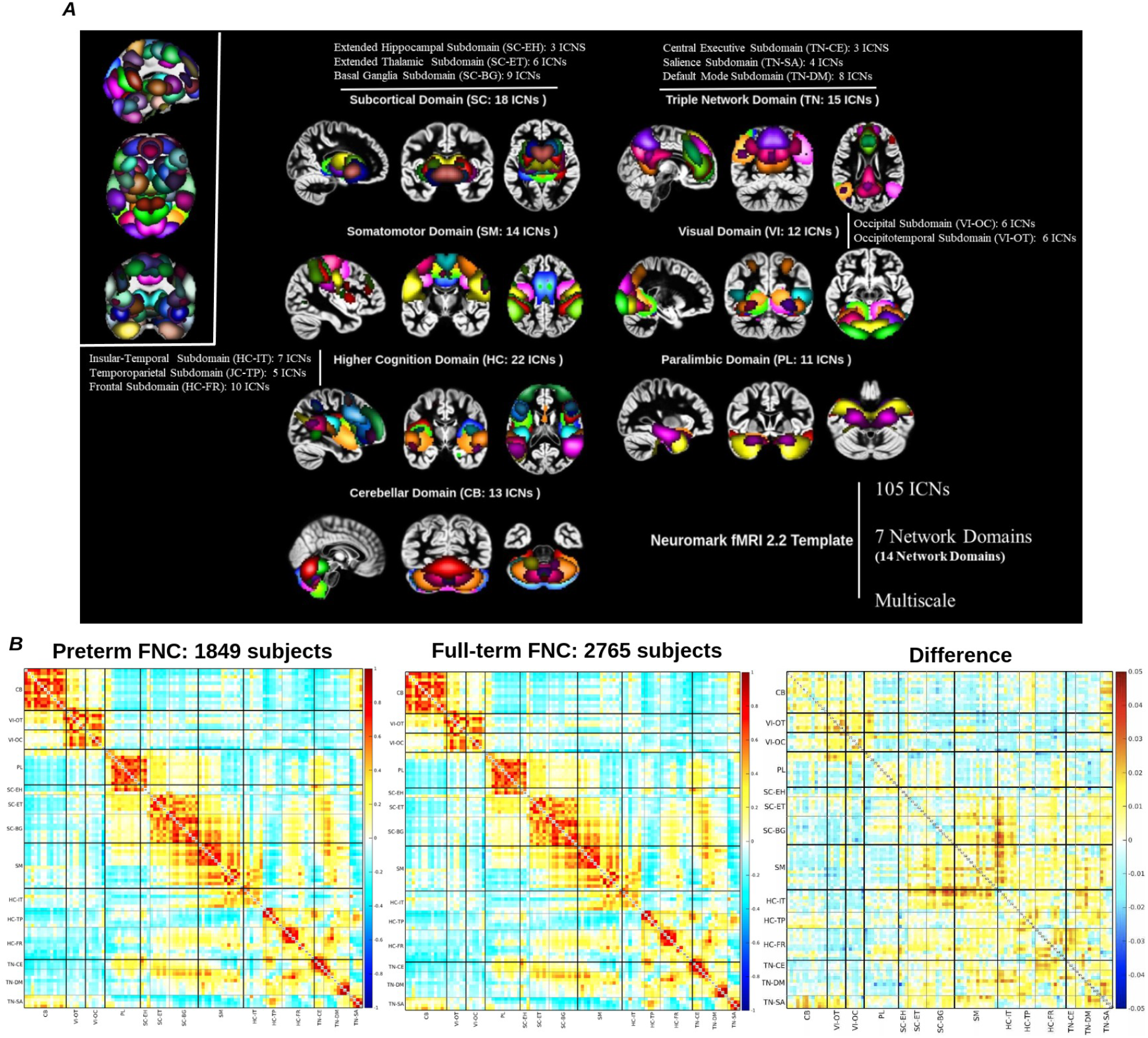
Preterm vs. Full-Term Multiscale Functional Connectivity Matrices. The multiscale brain network, NeuroMark2.2 Template (shown in ***A***), was used to extract multiscale time series from the ABCD dataset. This process generated subject-specific estimates of 105 intrinsic connectivity networks (ICNs) and their corresponding time courses in resting-state fMRI. These 105 ICNs were then grouped into seven major brain networks: visual (VI: VI-OT, VI-OC), cerebellar (CB), subcortical (SC: SC-EH, SC-ET, SC-BG), somatomotor (SM), paralimbic (PL), higher cognitive (HC: HC-IT, HC-TP, HC-FR), and the triple network (TN: TN-CE, TN-SA, TN-DM). The average preterm functional connectivity matrix, based on 1,849 subjects, and the average full-term functional connectivity matrix, based on 2,765 subjects, was shown in ***B***, along with the difference functional connectivity between preterm and full-term subjects.

Additionally, a two-sample t-test was performed on each element of the upper triangular portion of the functional connectivity matrix to compare preterm and full-term subjects. After applying False Discovery Rate (FDR) correction for multiple comparisons, 419 connections showed significant differences. This indicates that 419 pairs of functional connections exhibited significant differences in functional connectivity between the two groups. These identified significant connections were primarily found in the VI, SM, and HC regions, which align with what we observed in the residual functional connectivity map.

### B. Three Brain Network Energies Showed Significant Differences Between Preterm and Full-term Subjects

The functional network energies were measured at both the whole brain network and subnetwork levels, as shown in ***Fig.2***. First, using two-sample t-test and FDR correction, a significant difference in network energy was identified at the whole brain level (*p*_*corrected*_ = 0.004, t = -2.883) between preterm and full-term subjects. Then, at the subnetwork level, we identified significant differences in the following regions: VI-OT (*p*_*corrected*_ = 0.0002, t = -3.741), VI-OC (*p*_*corrected*_ = 0.013, t = -2.488), SM (*p*_*corrected*_ = 0.001, t = -3.216), HC-TP (*p*_*corrected*_ = 0.0002, t = -3.779), and HC-FR (*p*_*corrected*_ = 0.002, t = -3.054). In other subnetworks, we did not capture significant differences in network energy between preterm and full-term subjects. These identified subnetworks are with VI, SM, and HC networks, with VI-OT and HC-TP showing the strongest significance differences.

**Fig. 2:**
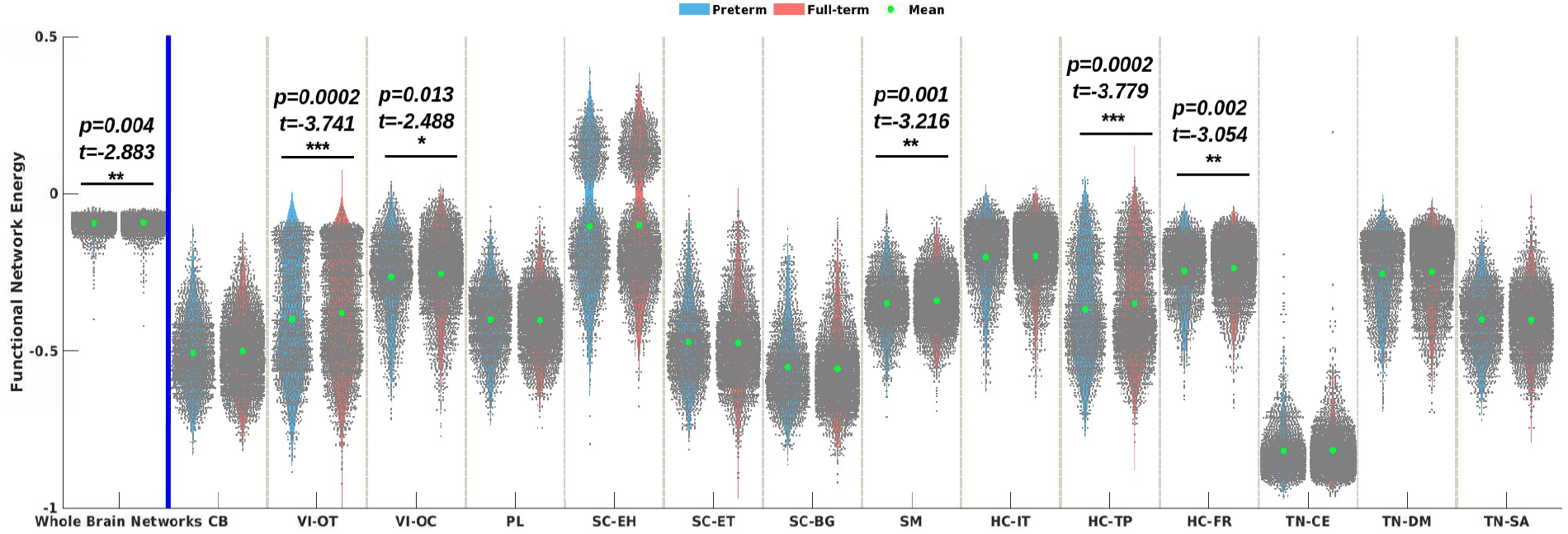
Functional Network Energy in the Whole-Brain Network and Its Subnets. The functional network energy was measured in the whole-brain network and in each subnetwork separately. The significant differences were observed specifically in the whole-brain network, as well as in the VI (VI-OT; VI-OC), SM, and HC (HC-FR; HC-TP) subnets (*, p<0.05; **, p<0.01; ***, p<0.001; FDR corrected.)

A further permutation test was performed to confirm that the above significant results were not caused by random chance. The results are presented in ***Fig.3***. As observed, the whole brain networks (*p*_*corrected*_ = 0.0026), VI-OC (*p*_*corrected*_ = 0.0118), VI-OT (*p*_*corrected*_ = 0.0003), SM (*p*_*corrected*_ = 0.0012), HC-FR (*p*_*corrected*_ = 0.0028), and HC-TP (*p*_*corrected*_ = 0.0003) showed significant differences, confirming that these differences were not due to random chance.

**Fig. 3:**
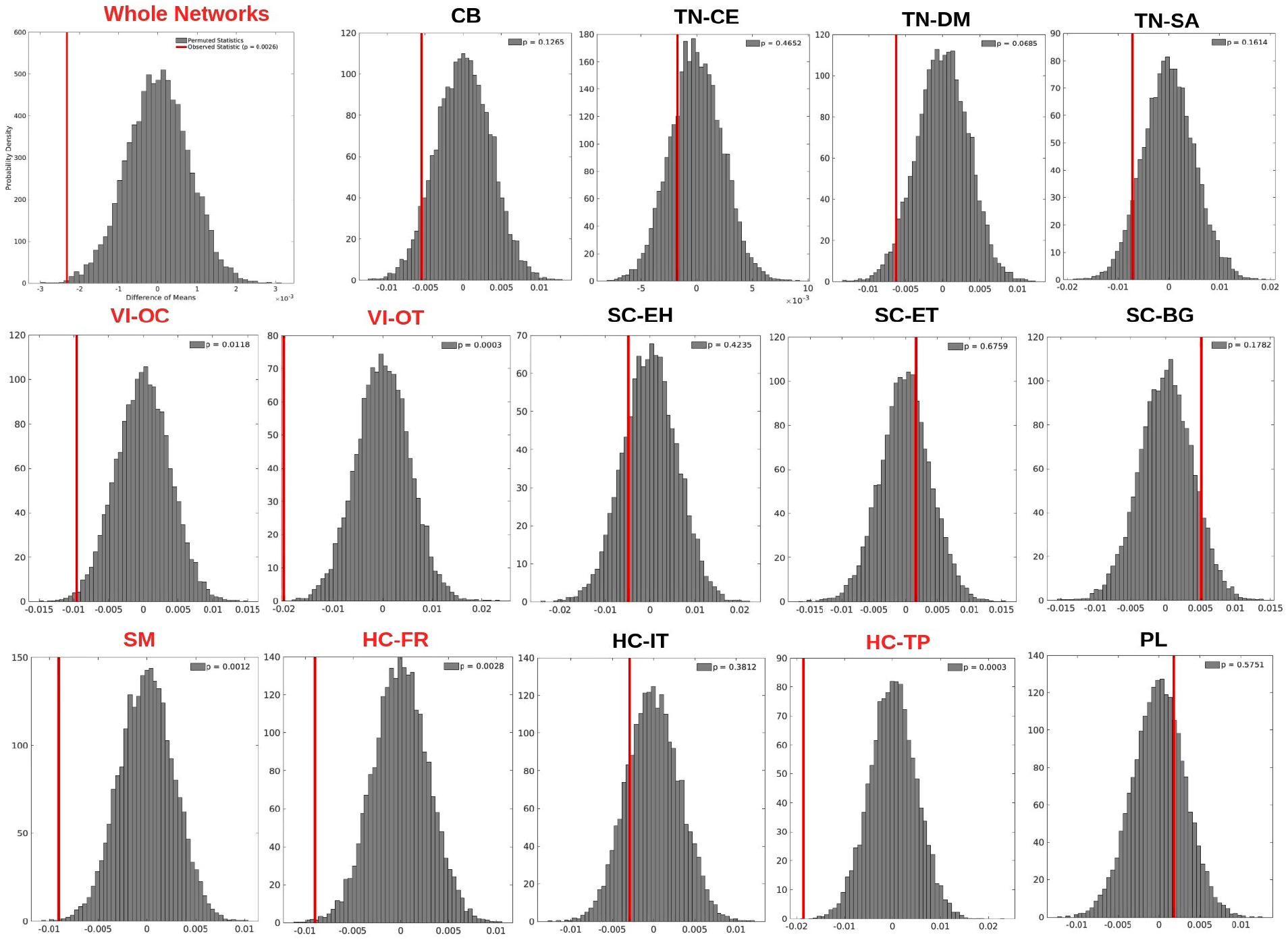
Permutation Test of Whole-Brain Network and Subnets. The permutation test was repeated 10,000 times for whole-brain network and subnetworks to assess whether there were significant differences in network energy statistics between preterm and full-term groups. The observed p-value (after FDR correction) was smaller than 0.05, indicating that the difference between the preterm and full-term groups was significant and not due to random chance. Significant whole-brain networks and subnetworks (VI-OC, VI-OT, SM, HC-FR, HC-TP) were highlighted in red.

### C. Full-term Subjects Exhibit Increased Flexibility across Three Key Canonical Functional Brain Networks Compared to Preterm Subjects

In both preterm and full-term subjects, the triadic network interactions are dominated by stable, balanced triangles. However, the brain networks of preterm subjects exhibit more stability, characterized by reduced energy levels. This might be related to difficulties with dynamic switching and adaptability in behavior [18]. In contrast, the networks of full-term subjects show greater instability, reflected in higher energy levels, which may facilitate more dynamic reconfiguration of information and enhance the brain’s flexibility [18].

These observed results can be explained using the Energy Landscape Hypothesis [23], [24], which suggests that the activity of the three identified canonical networks becomes too stable in preterm subjects, causing the networks to remain in a fixed state for longer periods. As shown in ***Fig.4***, the depths of the energy landscape basins for these networks become deeper, contributing to their increased stability.

**Fig. 4:**
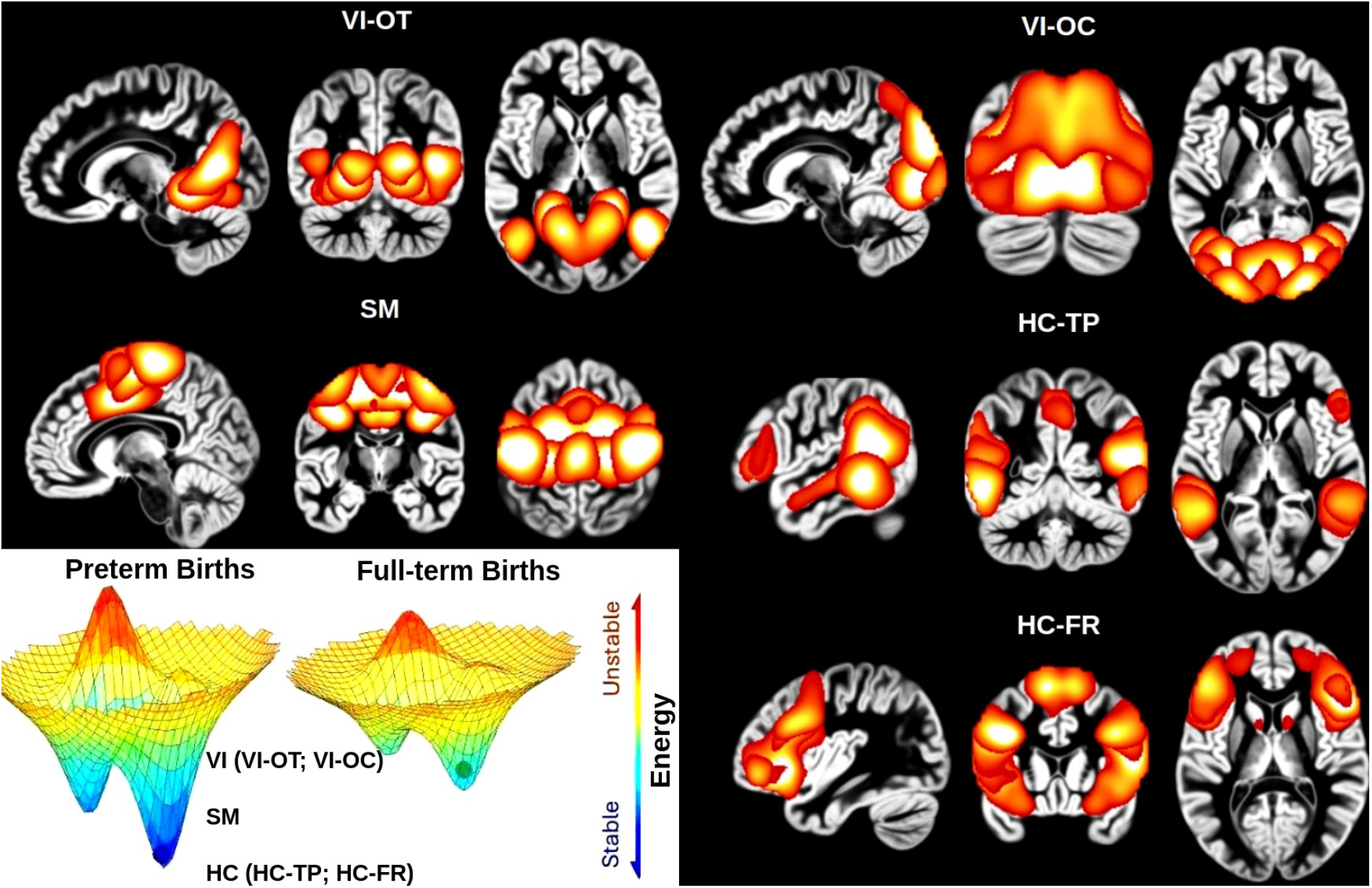
Spatial Maps of Key Networks and the Energy Landscape: A Comparison Between Preterm and Full-Term Subjects. The largest deviations in subnetworks between preterm and full-term groups in functional network energy were observed in VI-OT, VI-OC, SM, HC-TP, and HC-FR. The main differences were observed in the VI, SM, and HC subnetworks. The three key canonical brain networks identified were more stable in preterm subjects compared to full-term, with the depths of the energy landscape basins increasing. This increased stability, however, also reduces the dynamic reconfiguration and flexibility of brain information organization.

Overall, subjects who were full-term present higher energy levels compared to preterm subjects, both in the whole-brain network and several subnetworks (VI-OC, VI-OT, SM, HC-TP, and HC-FR), and the spatial maps of these three networks are illustrated in ***Fig.4***. This suggests a more complex and conflicting organization of functional connections, allowing for greater adaptability in response to cognitive demands.

## IV. Future Work

There are two main areas of focus for future work. First, we will examine specific brain components to identify key regions that play a major role in distinguishing preterm and full-term subjects neuronal development. Our study has identified significant differences in three key brain networks in 9-10 year old children that were born prematurely compared to their full-term peers. Within these networks, however, there are multiple components. The next goal is to isolate the specific components within these networks, enabling us to pinpoint the precise regions most affected by preterm birth. Second, once these regions are identified, we will expand our research to animal models, using rat brain images and models to guide us in understanding the underlying microscopic neural mechanisms that contribute to the developmental differences between preterm and full-term brains. This approach will provide a more comprehensive understanding of how preterm birth impacts brain development, ultimately offering valuable insights into the specific challenges faced by individuals who were born prematurely.

## V. Conclusions

In this study, we measured functional network energy between preterm and full-term subjects based on estimated multiscale functional connectivity from a subset adolescents from the ABCD study. We identified three key networks: visual, sensorimotor, and high cognitive, which show significant differences between preterm and full-term subjects. Additionally, we found that while preterm subjects exhibit low energy in these three key networks, they show reduced flexibility in brain information organization compared to full-term subjects, resulting in deficits within these networks in preterm subjects.

## Declaration of Competing Interest

The authors declare that they have no known competing financial interests or personal relationships that could have appeared to influence the work reported in this paper.

## ACKNOWLEDGMENT

This work was supported by NIH grant R01MH130595 and R34DA061483. The authors thank anonymous reviewers for constructive feedback for improving the manuscript.

## Notes

### Competing Interest Statement

The authors have declared no competing interest.

### Summary of Updates

Fixed some grammar and updated some content.

